# Intratumoral Delivered Novel Circular mRNA Encoding Cytokines for Immune Modulation and Cancer Therapy

**DOI:** 10.1101/2021.11.01.466725

**Authors:** Jiali Yang, Jiafeng Zhu, Yiyun Chen, Yaran Du, Yiling Tan, Linpeng Wu, Jiaojiao Sun, Mengting Zhai, Lixiang Wei, Na Li, Ke Huang, Qiangbo Hou, Zhenbo Tong, Andreas Bechthold, Zhenhua Sun, Chijian Zuo

**Affiliations:** Suzhou CureMed Biopharma Technology Co., Ltd. No. 388, Xinping street, Suzhou, 215000, China; Key Laboratory of Energy Thermal Conversion and Control of Ministry of Education, School of Energy and Environment, Southeast University, Nanjing, 210096, China; Institute of Pharmaceutical Biology and Biotechnology, Albert-Ludwigs-Universität Freiburg, Stefan-Meier-Straße 19, 79104 Freiburg, Germany

**Keywords:** Circular mRNA, IRES, cytokines, intratumoral injection, cancer therapy, tumor microenvironment

## Abstract

The application of mRNA as a novel kind of vaccine has been proved recently, due to the emergence use authorization (EUA) by FDA for the two COVID-19 mRNA vaccines developed by Moderna and BioNTech. Both of the two vaccines are based on canonical linear mRNA, and encapsulated by lipid nanoparticle (LNP). Circular mRNA, which is found to mediate potent and durable protein expression, is an emerging technology recently. Owing to its simplicity of manufacturing and superior performance of protein expression, circular mRNA is believed to be a disruptor for mRNA area. However, the application of circular mRNA is still at an initiation stage, proof of concept for its usage as future medicine or vaccine is necessary. In the current study, we established a novel kind of circular mRNA, termed C-RNA, based on Echovirus 29 (E29)-derived internal ribosome entry sites (IRES) and newly designed homology arms and RNA spacers. Our results demonstrated that this kind of circular mRNA is able to mediate strong and durable protein expression, compared to typical linear mRNA. Moreover, for the first time, our study demonstrated that direct intratumoral administration of C-RNA encoding a mixture of cytokines achieved successful modulation of intratumoral and systematic anti-tumor immune responses and finally leading to an enhancement of anti-PD-1 antibody-induced tumor repression in syngeneic mouse model. Additionally, after an optimization of the circular mRNA formulation, a significant improvement of C-RNA mediated protein expression was observed. With this optimized formulation, C-RNA induced enhanced anti-tumor effect via intratumoral administration and elicited significant activation of tumor-infiltrated total T cells and CD8^+^ T cells. Collectively, we established C-RNA, a novel circular mRNA platform, and demonstrated that it can be applied for direct intratumoral administration for cancer therapy.

## Introduction

The Emergence Use Authorization (EUA) of Moderna’s and BioNTech’s Covid-19 mRNA vaccines by FDA at the year 2020 proved that, as a novel class of vaccine, mRNA vaccine meets the requirement for vaccine development and clinical applications^1^. The BioNTech’s Covid-19 mRNA vaccine presented excellent protective rate of preventing infection from Covid-19, either in Phase III clinical trial or in the real world^2,3^. Additionally, due to the high titer of neutralizing antibody and activation of T-cell immunity, mRNA vaccine is proven to provide protection against Covid-19 variants^4^. According to the supreme performances of mRNA vaccine during Covid-19 pandemic, it is believed that mRNA technology may deeply change the paradigm of vaccines and therapeutics development.

To the best of our knowledge, most of the mRNA pipelines that being evaluated in clinical trials all over the world are based on typical linear mRNA, which can be generated by RNA polymerase-mediated in vitro transcription (IVT). Linear mRNA consists of a Cap structure, 5’ and 3’ untranslational regions (UTRs), protein encoding region and a polyA tail^5^. The 5’ cap0 or 5’ cap1 structure is generated via enzymatic reaction after IVT procedure or co-transcriptional incorporation of cap analog during IVT procedure^6,7^. This Capping reaction that replaces the pre-existing 5’-PPP structure reduces its immunogenicity and mediates strong promotion of mRNA translational activity by means of the tight interaction of 5’ Cap structure and elongation initiation factors^8^. PolyA tail that generated by polyA polymerase after IVT step, with a length about 200 to 250 bp, facilitates mRNA stability and finally boosts protein production^9^. 5’ and 3’ UTRs can be engineered to promote mRNA stability and protein expression^10^. To minimize the immunogenicity of mRNA, modified nucleosides can be incorporated into mRNA strand in the step of IVT^11^. The complicated structure of linear mRNA determines the complexity of its mRNA manufacturing and maybe a relative low yield after a few steps of synthesis and purifications.

In addition to linear mRNA, there is an emerging research area focusing on circular mRNA, which is proven to mediate potent and durable protein expression in vitro^12^. Despite the fact that most of the circular RNAs found in eukaryotic cells so far are considered as non-coding RNAs^13^, some circular RNAs are demonstrated to encode and express proteins^14^. Most of the circular RNAs that have been identified in eukaryotes are derived from in vivo back splicing mechanism^15^. After decades of investigations, methods for in vitro circularization of RNAs have been developed. In summary, circular RNAs can be generated in vitro through different methods^16^, including enzymatically ligation via T4 ligases, chemical strategies for the ligation of 5’ end and 3’ end, and finally a ribozyme strategy, by which an intra-molecular covalent bond is formed and circular RNAs is generated.

In 1992, M.Puttaraju and Michael D.Been reported that based on Anabaena group I intron, they invented a Group I permuted intron-exon (PIE) system to conduct an ribozyme catalyzed in vitro self-splicing and finally generate an exon-exon covalent -ligated circular RNA with high efficiency. In this case, an insertion of foreign nucleic acid sequences into the sites between exons and introns can generate a circular RNA that containing a foreign RNA fragment^17^. Based on this PIE system, R. Alexander Wesselhoeft et.al made optimizations by adding homologous arms and spacers to facilitate the generation of large-size circular RNAs and by adding internal ribosome entry site (IRES) element to mediate efficient protein translation^12^. They screened a diversity of viral-origin IRES elements and identified the IRES Coxsackievirus B3 (CVB3) as a high efficient one for cap-independent translation. Importantly, they found that this kind of CVB3 IRES-driven circular mRNA conducts stronger and more durable protein expression in vitro, compared to typical linear mRNA, with or without nucleoside modifications^12^. Their further studies indicated that circular mRNA after HPLC-purifications exhibited very low mRNA immunogenicity, suggesting that purified circular mRNA is probably suitable for applications in vivo as vaccines or therapeutic medicines^18^. However, more evidence is required to prove that circular mRNA is a type of proper molecule for clinical usage, especially *in vivo* experiments and clinical trials.

In the current study, we established a novel kind of circular mRNA, termed “C-RNA”, based on Echovirus 29 derived IRES and newly designed homology arms as well as RNA spacers. Compared to typical linear mRNA, C-RNA can mediate strong and durable protein expression. Moreover, for the first time, our study demonstrated that circular mRNA encoding a mixture of cytokines can be directly used for intratumoral administration to modulate intratumoral and systematic anti-tumor immune responses, finally lead to an enhancement of anti-PD-1 antibody induced tumor repression in syngeneic mouse models. Additionally, after an optimization of the circular mRNA formulation, a significant improvement of C-RNA-mediated protein expression *in vivo* was observed. With this optimized formulation, C-RNA induced better anti-tumor effect via intratumoral administration, and elicited significant activation of tumor-infiltrated CD4^+^ and CD8^+^ T cells. Collectively, we established C-RNA, a novel circular mRNA platform, and demonstrated that it can be applied for direct intratumoral administration for cancer therapy.

## Materials and Method

### Gene cloning and vector construction

DNA fragments that containing PIE elements, IRES, coding regions and others were chemically synthesized and cloned into a restriction digestion linearized pUC57 plasmid vector. The vector for linear mRNA containing beta-globin 5’ UTR and tandem beta-globin 3’ UTR, and coding regions was chemically synthesized and cloned into a restriction digestion linearized pUC57 plasmid. DNA synthesis and gene cloning were customized ordered from Suzhou Genwitz Co.Ltd. (Suzhou, China).

### circRNA preparations

circRNA precursors were synthesized by invitro transcription from a linearized plasmid DNA template using a Purescribe™ T7 High Yield RNA Synthesis Kit (CureMed, Suzhou, China). After in vitro transcription, reactions were treated with DNase I (CureMed, Suzhou, China) for 15 min. After DNaseI treatment, unmodified linear mRNA was column purified using a GeneJET RNA Purification Kit (Thermo Fisher). For circRNA: RNA was purified, after which GTP was added to a final concentration of 2 mM along with a buffer including magnesium (50 mM Tris-HCl, (pH 8.0), 10 mM MgCl2, 1 mM DTT; Thermo Fisher). RNA was then heated to 55 °C for 15 min, and then column purified. RNA was separated on 1% agarose gels using the agarose gel systems (Bio-Rad); ssRNA Ladder (Thermo Fisher) was used as a standard.

### circRNA purification

For high-performance liquid chromatography (HPLC), RNA was run through a 30 × 300 mm size exclusion column with particle size of 5 μm and pore size of 1000 □ (Sepax Technologies, Suzhou, China) on an SCG protein purification system (Sepure instruments, Suzhou, China). RNA was run in RNase-free Phosphate buffer (pH:6) at a flow rate of 15 mL/minute. RNA was detected and collected by UV absorbance at 260 nm. Concentrate the purified circRNA in an ultrafiltration tube, and then replace the phosphate buffer with an RNase-free water.

### Linear mRNA preparations

The plasmid vector was digested by XbaI for linearization, and transcription was carried out using HyperScribe™ All in One mRNA Synthesis Kit (APExBio), with Cap1 analog incorporation and N1-methyl-pseudouridine as modified nucleoside in transcription. RNA was purified by using a GeneJET RNA Purification Kit (Thermo Fisher).

### Cell culturing

HEK293T and the murine colon adenocarcinoma cell line MC38 as well as the murine melanoma cell line B16F10 were purchased from Cobier Biosciences (Nanjing, China) and cultured in Dulbecco’s modified Eagle’s medium (DMEM, BI) supplemented with 10% fetal calf serum (BI) and penicillin/streptomycin antibiotics (100 U/ml penicillin, 100μg/ml streptomycin; Gibco). The human non-small cell lung cancer cell line A549 and NCI-H358 were purchased from Cobier Biosciences (Nanjing, China) and cultured in Dulbecco’s modified Eagle’s medium (DMEM, BI) and RPMI 1640 (BI) respectively supplemented with 10% fetal calf serum (BI) and penicillin/streptomycin antibiotics (100 U/ml penicillin, 100μg/ml streptomycin; Gibco). All cells were maintained at 37 °C, 5% CO2, and 90% relative humidity.

### In vitro transfection of circRNA

For the circRNA transfection in HEK293T, 1×10^5^ cells per well were seeded in 24-well plates or 6-well plates. For 24-well plates, 500 ng circRNAs, or for 6-well plates, 2 μg circRNAs were transfected into each well of the cells using Lipofectamine MessengerMax (Invitrogen, LMRNA003) according to the manufacturer’s instructions. Cells were collected at 24 h post transfection for the following detections.

### Protein expression analysis

GFP fluorescence was observed and imaged using Olympus IX70 Microscope Cutaway Diagram 24 h after transfection. Average cell fluorescence intensity was detected by flow cytometry 24 h after transfection. HEK293T-GFP and HEK293T control cells were trypsinized and suspended in DMEM supplemented with 10% FBS and 1% penicillin/streptomycin. Cells were then washed twice and resuspended in PBS (Thermo Fisher) according to the manufacturer’s instructions. Fluorescence was detected for 10,000 events on a BD FACSCelesta™ flow cytometer (BD Biosciences).

### Encapsulation of circular mRNA by lipid nanoparticle (LNP)

Circular mRNA-LNP complex was generated through microfluidic devices (Micro&Nano Technologies, Shanghai, China). Purified circular mRNA was dissolved in citric acid buffer and lipids were dissolved in ethanol. The liquid flow rate was set as 12 mL/min, mRNA/lipids (v/v) was set as 3:1. Circular mRNA-LNP complex was purified by filtration.

### Mice

C57BL/6 female mice and CD-1 nude female mice were purchased from Charles River (Beijing, China) at the age of 6-8 weeks and housed in specific-pathogen-free facilities and all experiments were conducted in accordance with procedures approved by the Institutional Animal Care and Use Committee (IACUC).

### In vivo tumor experiments

For tumor implantation, mice were injected with A549, NCI-H358, B16F10 or MC38 (1×106 cells per animal) s.c. in the right flank. Tumor growth was monitored by 2 perpendicular diameters with a digital caliper every 2 to 4 days and tumor volume was calculated by the modified ellipsoidal formula: V = ½ (Length × Width^2^). Intratumoral injections were performed with 1 mL 29G×1/2 insulin syringe (BD).

### Bioluminescence imaging

Mice were anesthetized in a chamber with 2.5% isoflurane and intra-peritoneal injections of D-luciferin (150mg/kg, Promega). Bioluminescence was measured by an IVIS Spectrum imaging system (PerkinElmer) while maintaining 2.5% isoflurane in the imaging chamber via a nose cone. Images were captured 10 minutes after luciferin administration at indicated time points. The photon flux values(photons/second), corresponding to the ROI (region of interest) marked around the bioluminescence signal, were analyzed using Living IMAGE software.

### Tumor samples preparation and flow cytometry

Tumors were isolated and dissected into around 1 mm3 pieces. Then these pieces were incubated in DMEM medium containing 1mg/mL collagenase-IV (Thermo Fisher) and 100 U/mL DNase-I (Novoprotein) at 37 °C for 30 minutes with gently shaking. Dissociated cells were filtered by a 70-μm nylon mesh filter to obtain single cell suspensions. For antibody staining,cells were firstly stained with eBioscience™ Fixable Viability Dye eFluor™ 780 (1:1000, Thermo Fisher) and then washed twice with PBS containing 2% FBS and 2 mM EDTA. Afterword, cells were stained with the following antibodies (1:100 in PBS containing 2% FBS and 2 mM EDTA) : APC anti-mouse CD3ε Antibody (Biolegend, cat 152306) , Brilliant Violet 605™ anti-mouse CD4 Antibody (Biolegend,cat 100451),Brilliant Violet 421 anti-mouse CD8a (Biolegend,cat 100737),FITC anti-mouse CD69 Antibody (Biolegend, cat 104506),PE anti-mouse CD45 (Biolegend,cat 103106). After incubating for 20 minutes in the refrigerator, cells were washed twice with PBS containing 2% FBS and 2mM EDTA, and analyzed by flow cytometry (Life,Attune NXT).

### PBMC isolation and in vitro studies

Peripheral blood samples (buffy coats) from healthy volunteers were isolated by Ficoll Hypaque gradient separation (Ficoll-Paque-Plus) and washed 3 times with PBS supplemented with 1 mM EDTA. Freshly isolated PBMC seeded at 96-well plate with the density of 500000/well and treated with cytokine mixture which were collected from different mRNA transfected 293T supernatant for 24 hours. And then the IFN-gamma production was determined by Human IFN-gamma ELISA kit (Neobioscience).

### Statistical Analyses

Two-tailed Student t test was used to determine statistical significance for data comparisons at a single time point. Two-way ANOVA was used to determine statistical significance for data comparisons with multiple time points. Mantel–Cox log rank test was performed to determine statistical significance for the comparison of survival curves. Prism version 8.0 (GraphPad) was used for generation of all graphs and performance of statistical analyses. Statistical significance shown for survival curves represents a comparison of the 2 survival curves. Statistical significance is denoted as ns, not significant, *P < 0.05, **P < 0.01, and ***P < 0.001.

## Results

### Construction and confirmation of C-RNA, a novel circular mRNA format

Due to the fact that CVB3 IRES derived circular mRNA presents high activity for protein translation, we sought to investigate alternative virus-origin IRES that can mediate high and durable protein expression in mammalian cells. In our study, the circular mRNA was generated by a vector the consists of a T7 promoter, PIE system, IRES and novel RNA spacers. After a transcription reaction mediated by T7 RNA polymerase, the RNA precursor was produced. Next, circular mRNA was generated after a further circularization procedure. As shown in Fig.1A and 1B, our results indicated that, a series of echovirus originated IRES exhibited significant activity of directing circular RNA translation in 293T cell line. IRES echovirus 24 (E24), echovirus 29 (E29) and echovirus 33 (E33) were found to direct moderate or strong EGFP expression. Among them, E29 was the preferable one. Furthermore, we designed chimeric IRES components in which the CVB3 core structure was inserted into an echovirus IRES basic framework. CVB3 core structure is defined as the RNA domain that interacts with RNA translation complex. However, our results showed that the chimeric IRES didn’t present significant improvement of protein expression (Fig.1A and 1B). E29 IRES and novel spacers based circular mRNA mediated the expression of EGFP last to even eleven days in HEK-293T (data not shown). Thus, IRES E29 was designated as the supreme IRES component for our circular mRNA platform. Collectively, our novel circular mRNA format, which is termed C-RNA, consists of E29 IRES, the protein coding region, spacers and Exon2-Exon1 junction that derived from PIE system (Fig.1G). Besides, we evaluated the expression of C-RNA with or without HPLC-grade purification in A549 cell line 24 h after C-RNA transfection. We found that the purified C-RNA mediated remarkable higher protein expression in A549, compared to the unpurified C-RNA (Fig.1E and 1F). The promotion of protein expression may cause by the removal of immunogenic 5’-ppp small intron fragments that generated after circularization reaction. These results suggested that purification by HPLC is a critical step with regards to C-RNA, no matter for elimination of immunogenicity or for promotion of protein expression.

**Fig.1.**
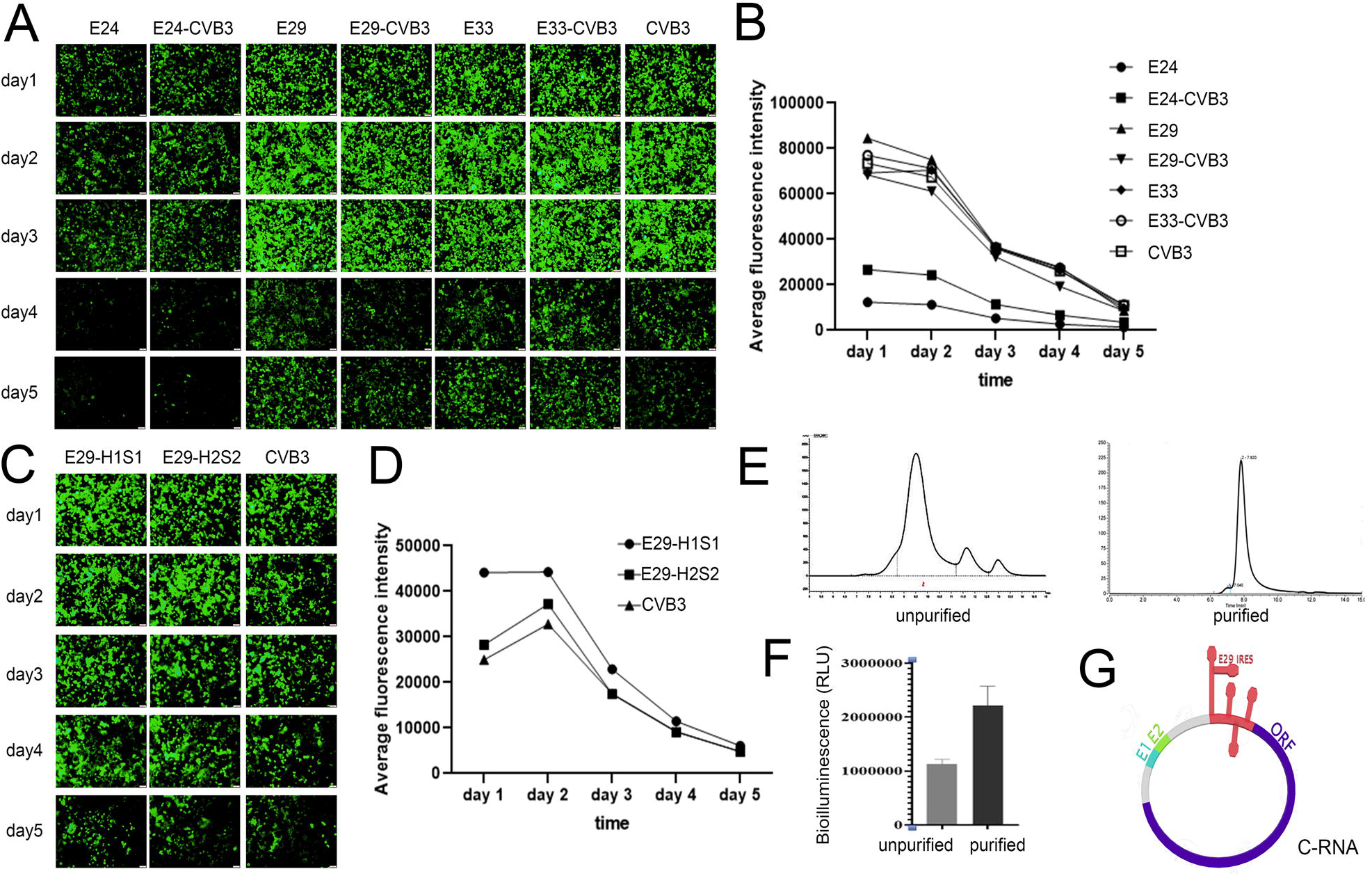
(A). Series of IRES derived from echovirus were tested in circular mRNA for measuring their activity of EGFP translation. Picture shows the expression of EGFP during day1 to day5; (B) Quantification of Fig.1A by FACS; (C) EGFP protein expression of circular mRNAs with combinations of IRES E29, homologous arms and spacers were examined during day1 to day5; (D) Quantification of Fig.1C by FACS; (E) Diagram of HPLC shows the elution of circular mRNA before (left) and after (right) HPLC-SEC purifications; (F) Luciferase activities were measured 24 h post transfection of C-RNA with or without HPLC-SEC purifications; (G) Scheme of C-RNA.

### C-RNA directed protein expression in vitro and in vivo

Next, we sought to investigate whether C-RNA is able to direct the expression of different types of proteins in vitro or in vivo. To delineate the localization of in vivo protein expression, C-RNA encoding luciferase encapsulated by lipid nanoparticle (LNP) was delivered to mice, and the expression of luciferase bioluminescence was detected by optical in vivo imaging system. As shown in Fig.2A, our results indicated that 6h and 24 h after i.m. injection, potent luciferase activity was found in the area of mouse liver and the injection site of muscle, suggesting that LNP delivered C-RNA majorly to liver and C-RNA mediated high level of protein expression. Importantly, the expression of luciferase can last to even 9 days in the injection site of muscle, suggesting that C-RNA mediated a durable protein expression in vivo. In addition to intracellular proteins (e.g., EGFP and luciferase), secreted proteins and transmembrane proteins were evaluated, and the in vivo protein expressions were examined. Firstly,to confirm C-RNA mediated transmembrane protein expression, we evaluated the in vitro expression of IL-15 CD8α and IL-15 GPI, two engineering transmembrane cytokines. Data shown in Fig.2B, at 24 h post RNA transfection, IL-15 CD8α and IL-15 GPI were found to express notably on cell membrane as detected by flow cytometry, suggesting that C-RNA mediated high and robust expression of these two transmembrane proteins. Next, if the expression of a secreted antigen trimer, receptor binding domain (RBD) of SARS-Cov2 was able to be directed by C-RNA was evaluated through in vitro transfection. As shown in Fig.2C, 24 h post the transfection of RBD C-RNA to 293T cell line, a high level of the secreted recombinant RBD trimer protein in the cell culture supernatant was detected by ELISA. To confirm whether C-RNA can mediate the secreted RBD protein expression in vivo, the circularized-and HPLC-purified RBD C-RNA was encapsulated by LNP and delivered to mouse. 24 h after intramuscular (i.m) injection of the C-RNA LNP complex, mouse serum was collected and significant level of RBD trimer protein was detected, suggesting that C-RNA mediated successful RBD protein expression in vivo. In summary, C-RNA can direct the expression of intracellular, secreted and transmembrane proteins in vitro. Additionally, the in vivo expression of intracellular protein and secreted protein were verified.

**Fig.2.**
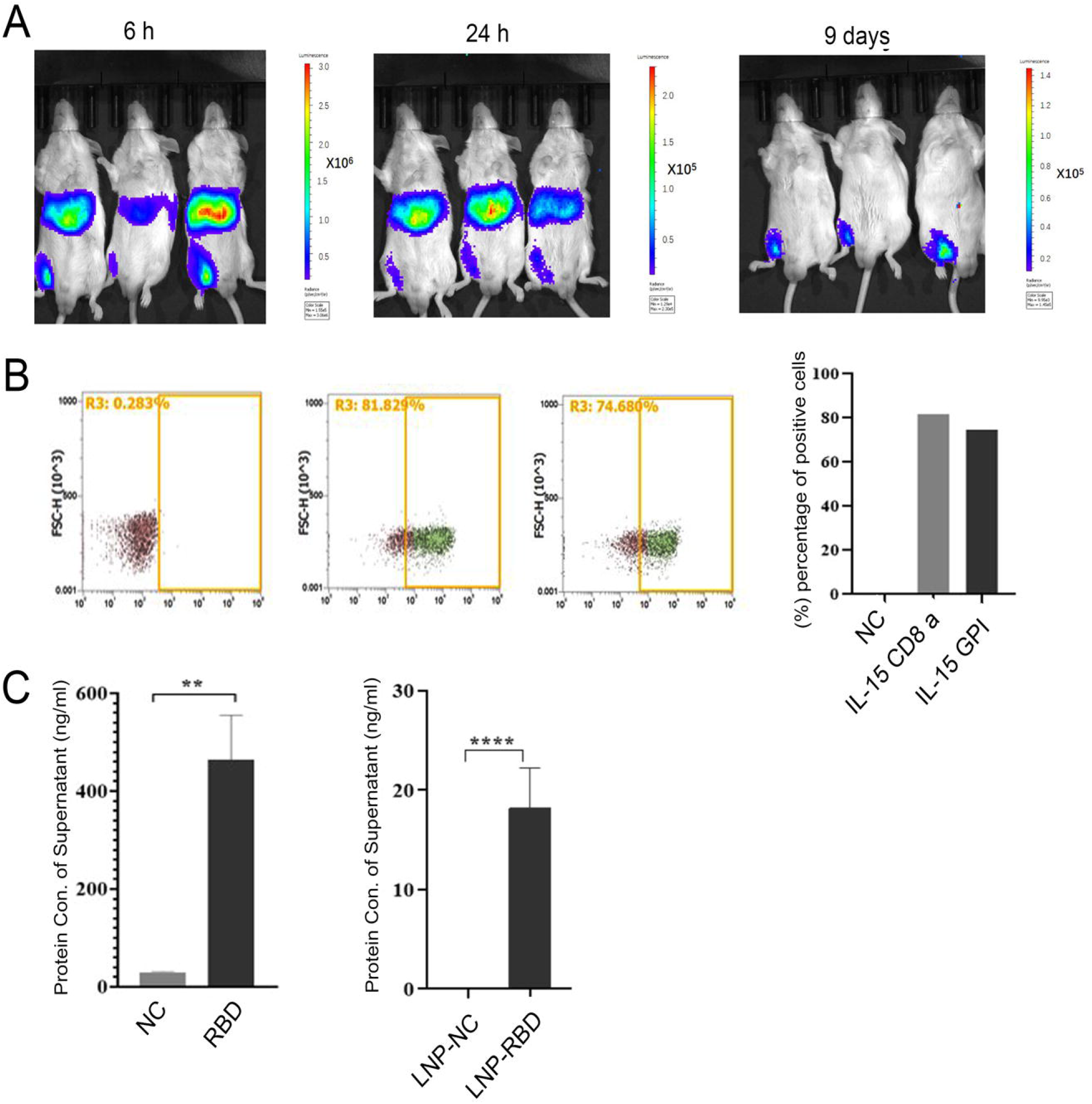
(A) Pictures show the bioluminescence of firefly luciferase at 6h, 24h and 9days after intramuscular injection of 10 μg LNP-C-RNA complex; (B) Detections of membrane protein IL-15 CD8 α and IL-15 GPI expression by FACS; (C) ELISA detections of the secreted form of RBD antigen in cell culturing supernatant of 293T at 24 h post 0.5 μg C-RNA transfection, and in mouse serum at 24 h post intramuscular injection of 10 μg LNP-C-RNA complex. **P<0.01, ****P<0.0001.

### Naked C-RNA can be directly delivered and specifically expressed in tumor tissue by intratumoral administration

In 2017, Yanping Kong *et*.*al* reported that cancer cells can spontaneously uptake DNA fragments with sizes ranging from 57 to 1620 bp, probably due to the over-activated endocytosis of cancer cells^19^. Based on this research, we proposed that circular mRNA may be uptaken by cancer cells in a similar way, despite that circular mRNA has a distinct shape and bigger size compared to the linear DNA. To confirm this hypothesis, C-RNA was used for direct intratumoral injection. C-RNA that encoding luciferase dissolved in PBS solution was injected into B16F10 xenograft tumor in C57BL/6, or injected subcutaneously, intradermally, intramuscularly, intraperitoneally, and intravenously. Our results indicated that significant bioluminescence was found specifically in B16F10 tumor after 6 h of C-RNA intratumoral injection (Fig.3A). However, there was no bioluminescence detected in the groups that with other varies injection methods, suggesting that C-RNA can be direct delivered and expressed specifically in tumor tissue. According to the report from Yanping Kong *et*.*al*, the direct delivery of naked DNA into cancer cells depends on characteristics of cancer cells. Thus, we broadened the types of xenograft tumors, including A549 and NCI-H358, cells derived from non-small-cell lung carcinoma, transplanted in nude mice; and MC38, cells derived from colon adenocarcinoma, transplanted in mice. Our results presented that 6h after intratumoral injection of C-RNA to all of these transplanted tumors yield significant bioluminescence in the tumor sites, suggesting that C-RNA was delivered and expressed in the tumor tissue, including A549, NCI-H358 and MC38 (Fig.3B), thus C-RNA shows a promising ability to express in varied kinds of tumors.

**Fig.3.**
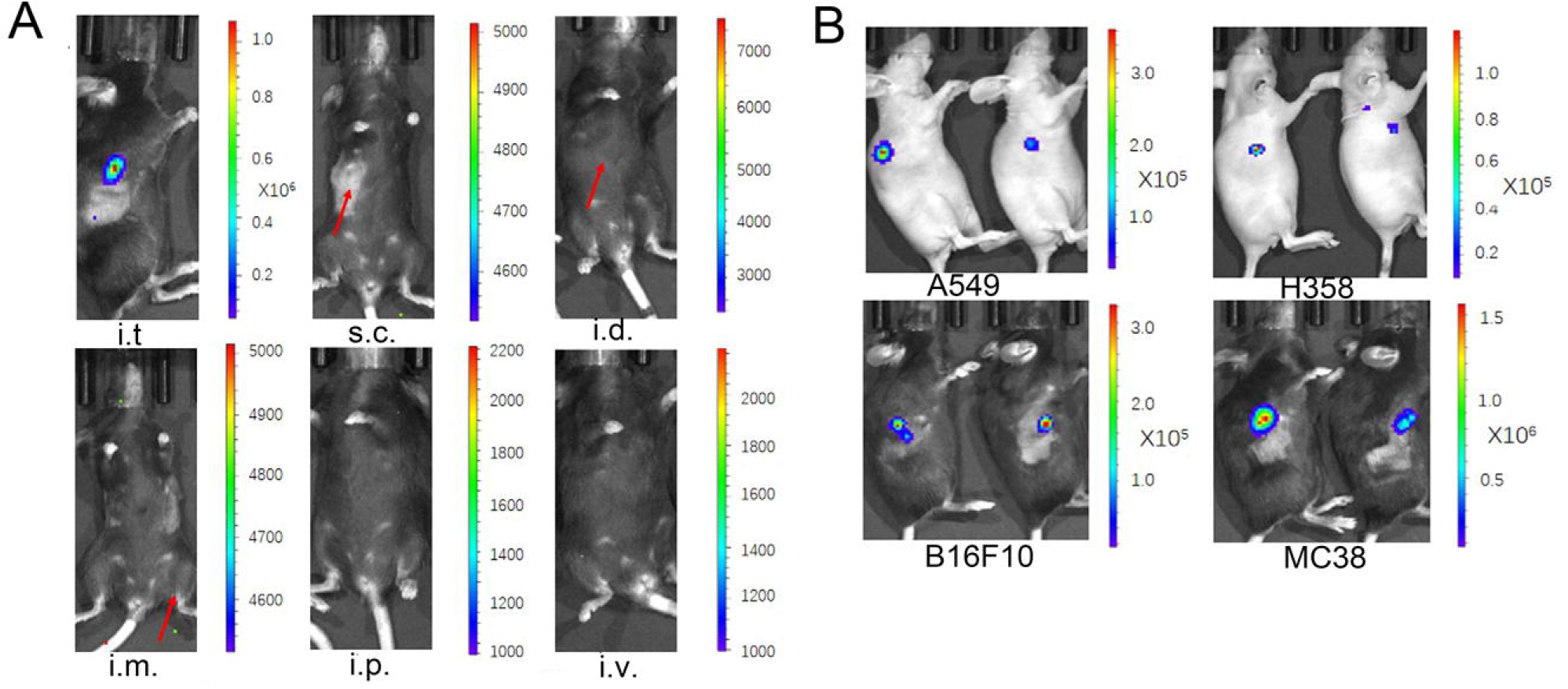
(A) C57BL/6 mice bearing B16F10 transplanted tumors were injected with naked 10 μg C-RNA-luciferase via indicated routes of administration. 6 hours later, images of bioluminescence measurement were taken. Scales represent light intensity in photons/sec; (B) CD-1 nude mice bearing A549 or H358 transplanted tumors and C57BL/6 mice bearing B16F10 or MC38 transplanted tumors were intratumorally injected with naked 10 μg naked C-RNA-lucifrease respectively. 6 hours later, images of bioluminescence measurement were taken. Scales represent light intensity in photons/sec.

### Intratumoral C-RNA presented higher and more durable protein expression than linear mRNA

Firstly, protein expressions of EGFP by C-RNA and typical linear mRNA were compared in 293T. As depicted by the in vitro study in Fig4A, C-RNA mediated higher and more durable protein expression than the typical linear mRNA in 293T cell line. Next, it still needs to be elucidated whether this phenomenon can be repeated in vivo, especially for the expression of C-RNA in tumor tissues. Thereof, the intratumoral luciferase expression directed by C-RNA and linear mRNA were compared head-to-head. The linear mRNA was constructed by typical beta-globin 5’ UTR and a tandem beta-globin 3’ UTR, cap1 cap structure and polyA structure derived from polyA polymerase. Intratumoral injections of naked C-RNA or linear mRNA both with the same amount and encoding the same *luciferase* gene into MC38 tumor in C57BL/6 were carried out, and bioluminescence measurements were taken at indicated time points. Results of the luciferase activities indicated that the expression of luciferase can be detected as early as 4 h, and notably the bioluminescent level of C-RNA group was higher than the linear mRNA group (Fig.4B and 4C). At the time points 4, 24, 48, 72 and 96 h, the bioluminescent levels were traced, and the results indicated that bioluminescent activities produced by C-RNA were about two folds higher than the linear one. Additionally, the bioluminescence level declined more slowly in C-RNA group, suggesting that C-RNA mediated higher and more durable protein expression in tumor tissue. These results are similar as the in vitro expression of C-RNA in 293T cell line.

**Fig.4.**
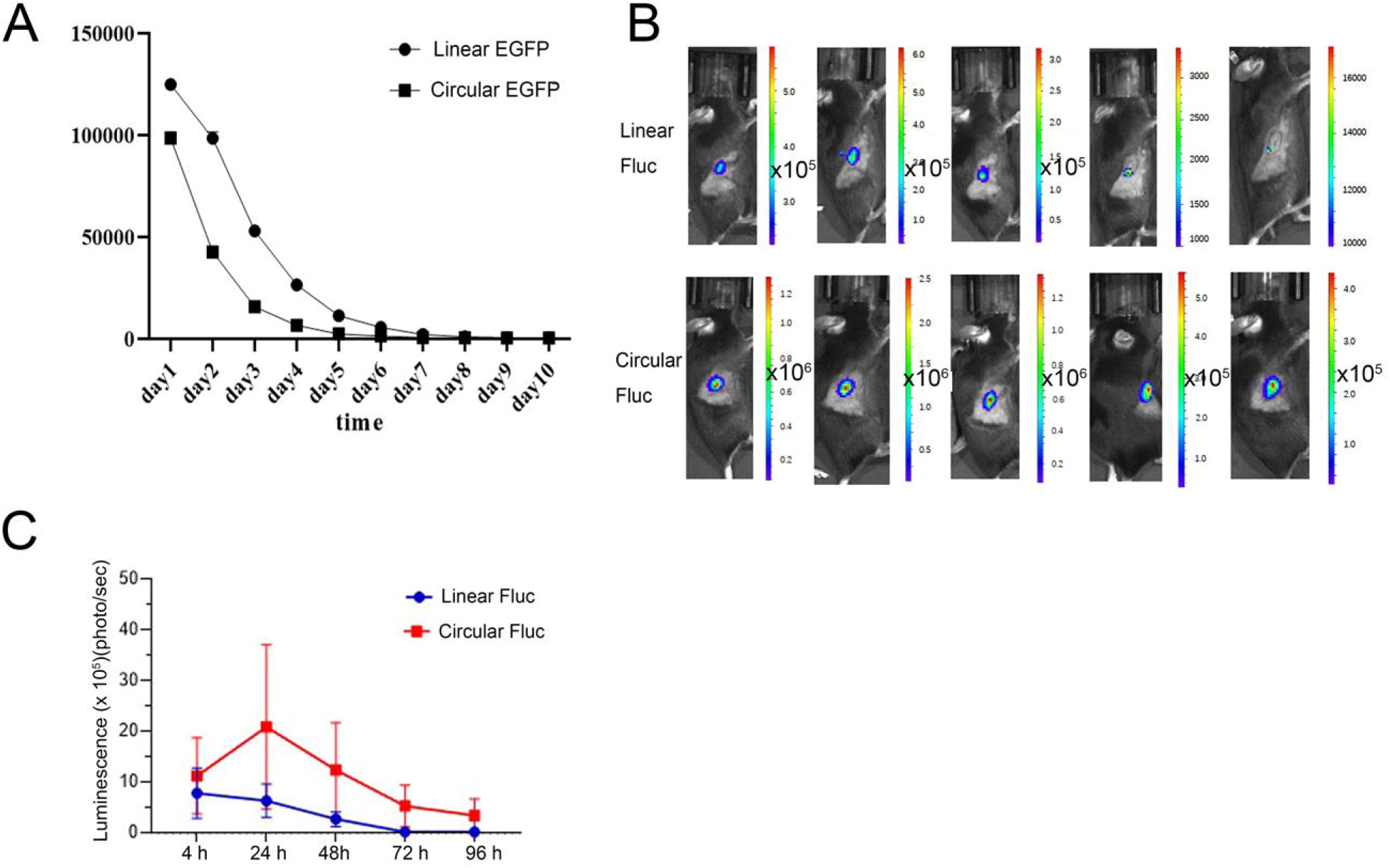
(A) C57BL/6 mice bearing MC38 transplanted tumors were intratumorally injected with 10 μg nucleotide modified linear luciferase mRNA (Linear Fluc) or C-RNA luciferase mRNA (Circular Fluc). Images of bioluminescence measurement were taken following indicated time. (B) Quantification of bioluminescence over time.

### Optimization of C-RNA formulation promoted intratumoral delivery and protein expression

In 2007, J Probst et.al reported that Ca^2+^ affected the delivery of naked linear mRNA into cells via local skin injection^20^. The optimization of mRNA solvent compositions led to a promotion of about 10-fold of linear mRNA uptake and protein expression^20^. It is still unknown how the intratumoral delivery of naked circular mRNA can be improved by adjusting the solvent compositions. In this study, different compositions were used as C-RNA solvent, including 0.9% NaCl, PBS, 1 x TE buffer and Ringer’s solution. Bioluminescence was measured 6 h later after C-RNA intratumoral administration to B16F10 xenograft tumors. Results indicated that C-RNA dissolved in Ringer’s solution brought to supreme luciferase expression in tumor sites (Fig.5A and 5B). According to the study from J Probst^20^, the compositions included in Ringer’s solution, such as CaCl2, NaCl, KCl and NaHCO3 may be vital for mRNA delivery. Thus, removal of the four compositions above in C-RNA solution was performed respectively, and again intratumoral injection to B16F10 tumors. We found that the removal of CaCl2 and KCl greatly reduced the bioluminescence level of C-RNA injection, and the removal of NaCl and NaHCO3 led to moderate and mild reduction, respectively (Fig.5A and 5B). These results suggest that CaCl2 and KCl are the two essential components for C-RNA intratumoral delivery.

**Fig.5.**
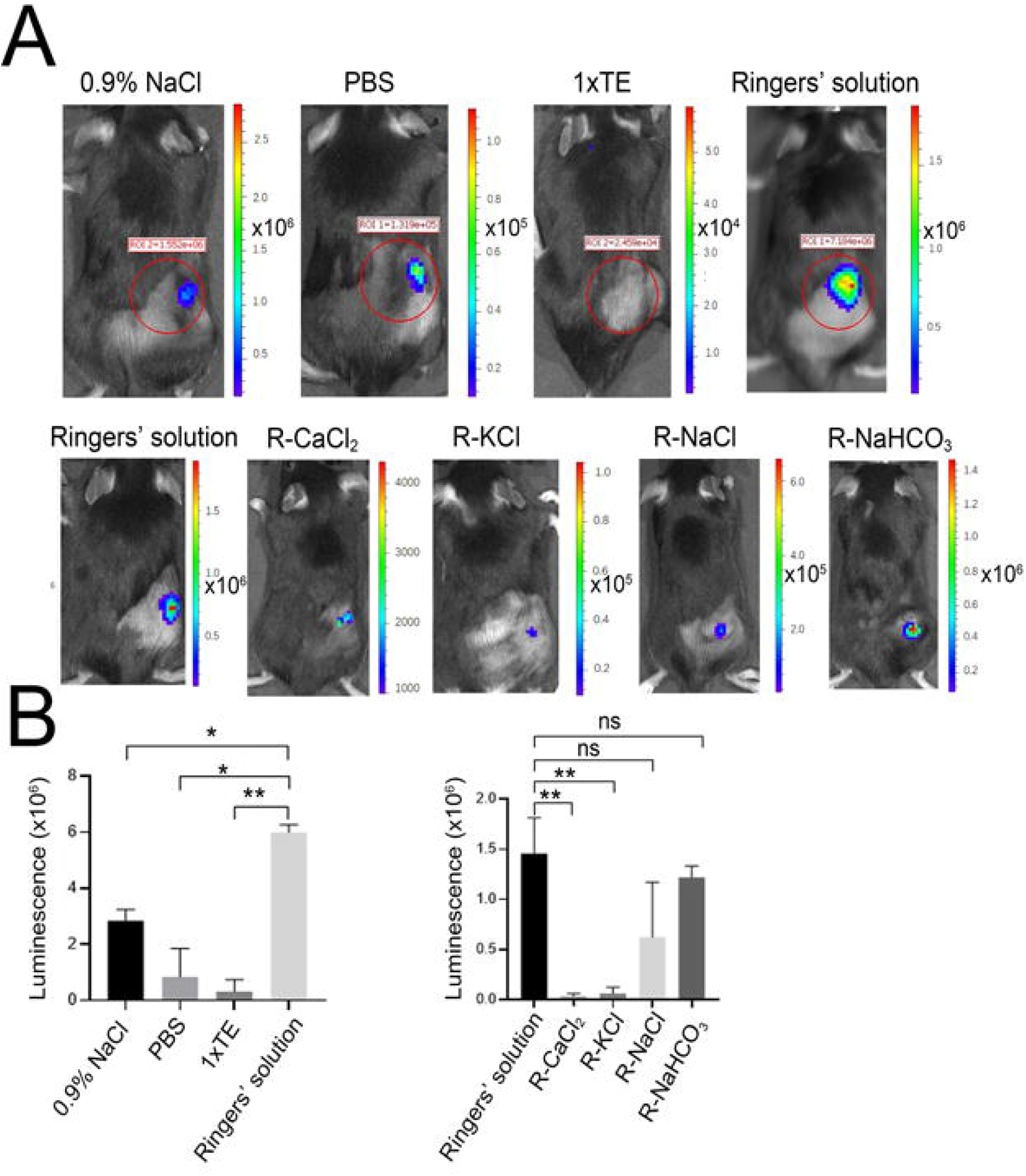
(A) Representative image of luciferase expression encoded by C-RNA dissolved in different solutions and in Ringers’ solution with removal of specific components. (B) Quantification of luciferase expression of (A).

### C-RNA encoding cytokines facilitated anti-PD-1 antibody mediated tumor suppression

Previous reports indicated that cytokines play critical roles in the regulation of tumor microenvironment. In this section of study, four cytokines including interleukin-15 (IL-15), interleukin-12 (IL-12) ,granulocyte macrophage colony-stimulating factor (GM-CSF) and interferon-alpha 2b (IFN-α 2b) were encoded by C-RNA. After transfecting C-RNAs that encoding four cytokines into 293T cells, the expression of secreted cytokines was quantified by ELISA (Fig.6A). High expression of these cytokines was found 24 h after transfection (Fig.6B). Furthermore, the cell supernatant of each transfection was collected and transferred to PBMC to assess their activities. We found that the incubation with each secreted cytokine resulted in remarkable up-regulation of CD69 positive CD4^+^ and CD8^+^ T cells in PBMC (Fig.6C), suggesting that the individual treatment of each cytokine led to activation of CD4^+^ and CD8^+^ T cells in vitro. Supreme activations were observed in the group treated with the mixture of secreted cytokines (Fig.6C). Next, C-RNA was dissolved in PBS or Ringers’ solution and directly delivered to tumors via intratumoral administration to modulate the tumor microenvironment. A mixture containing 10 μg of each C-RNA was injected intratumorally every three days, with or without i.p. administrated anti-PD-1 antibody. For B16F10 tumor, until 19 days of tumor inoculation, mice treated with cytokine C-RNA mix with PBS solvent, combined with anti-PD-1 antibody exhibited significant suppressed tumor growth, compared to the group with anti-PD-1 antibody monotherapy (data not shown). However, mice with combination treatment presented no significant improvement for the final survival rate in the 40-day survival tests, compared to antibody monotherapy group or negative control (data not shown). These results suggest that mice with B16F10 tumor responded to the combination treatment with C-RNA mixture and anti-PD-1 antibody. Next, the effect of C-RNA mixture with Ringers’ solution as solvent for the treatment of mice bearing B16F10 tumor was evaluated. After the same administration format, until 24 days of tumor inoculation, mice with combination treatment presented significant tumor growth inhibition (Fig.7A). Moreover, in the 50-day survival tests, a notable improvement of final survival rate was found after combination treatment, compared to negative control or antibody monotherapy, as shown in Fig.7B. These results suggest that mice bearing B16F10 tumor responded well to the combination treatment, leading to a significant tumor suppressive effect, finally resulted in improved survival rate. Additionally, we evaluated the anti-tumor effect of C-RNA mixture in the treatment of mice bearing MC-38 tumor. Intratumoral administration of C-RNA mixture with Ringers’ solution as solvent to MC-38 tumor resulted in notable retardation of tumor growth, either C-RNA monotherapy or combination with anti-PD-1 antibody. Combination treatment of C-RNA and anti-PD-1 antibody presented better control of tumor growth, suggesting that a coordination between cytokine mixture and anti-PD-1 antibody exists (Fig.7C). Importantly, in the 50-day survival tests, a notable improvement of final survival rate was found after C-RNA mixture monotherapy of combination treatment, featuring with 3/8 complete response (CR) rate, compared to negative control or antibody monotherapy, as shown in Fig.7D. In summary, these results suggest that cytokines-encoding C-RNA mixture exhibits supreme anti-tumor effect, especially combined with checkpoint blockade agents for tumor therapy.

**Fig.6.**
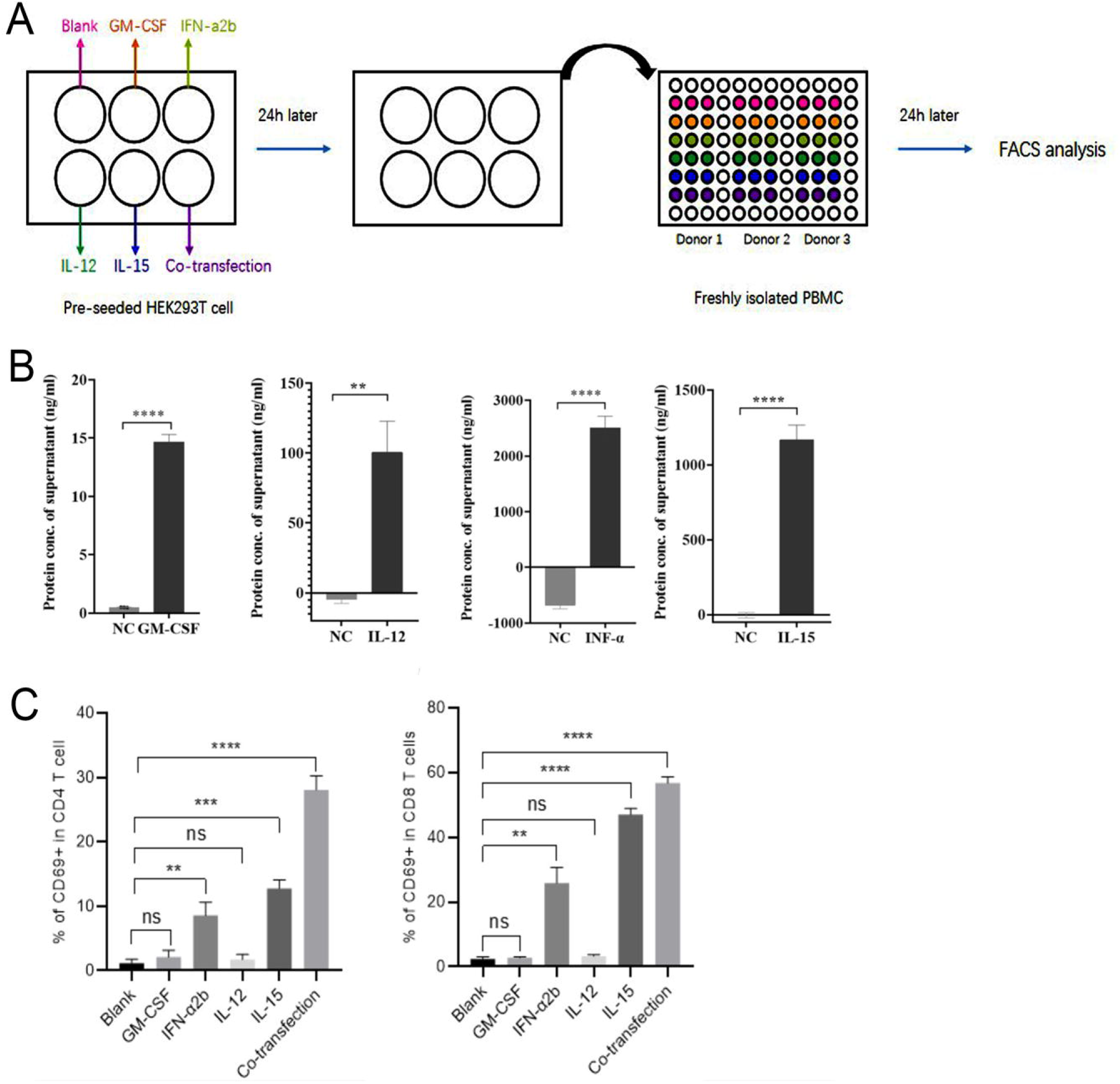
(A) Schematic representation of the experimental design. Pre-seeded HEK293T were transfected with indicated C-RNA. 24 hours later, the supernatant of each well was collected and centrifuged to remove 293T cells. PBMC isolated from three healthy donors were seeded in 96-well plate and then resuspended with 100 μL of indicated supernatant. After 24 hours of incubation, PBMC were stained with fluorophore-conjugated antibodies and analyzed by flow cytometry; (B) Percentage of CD69 positive CD4 T cells in total CD4 T cells, where each spot represents one blood donor; and percentage of CD69 positive CD8 T cells in total CD8 T cells, where each spot represents one blood donor.

**Fig.7.**
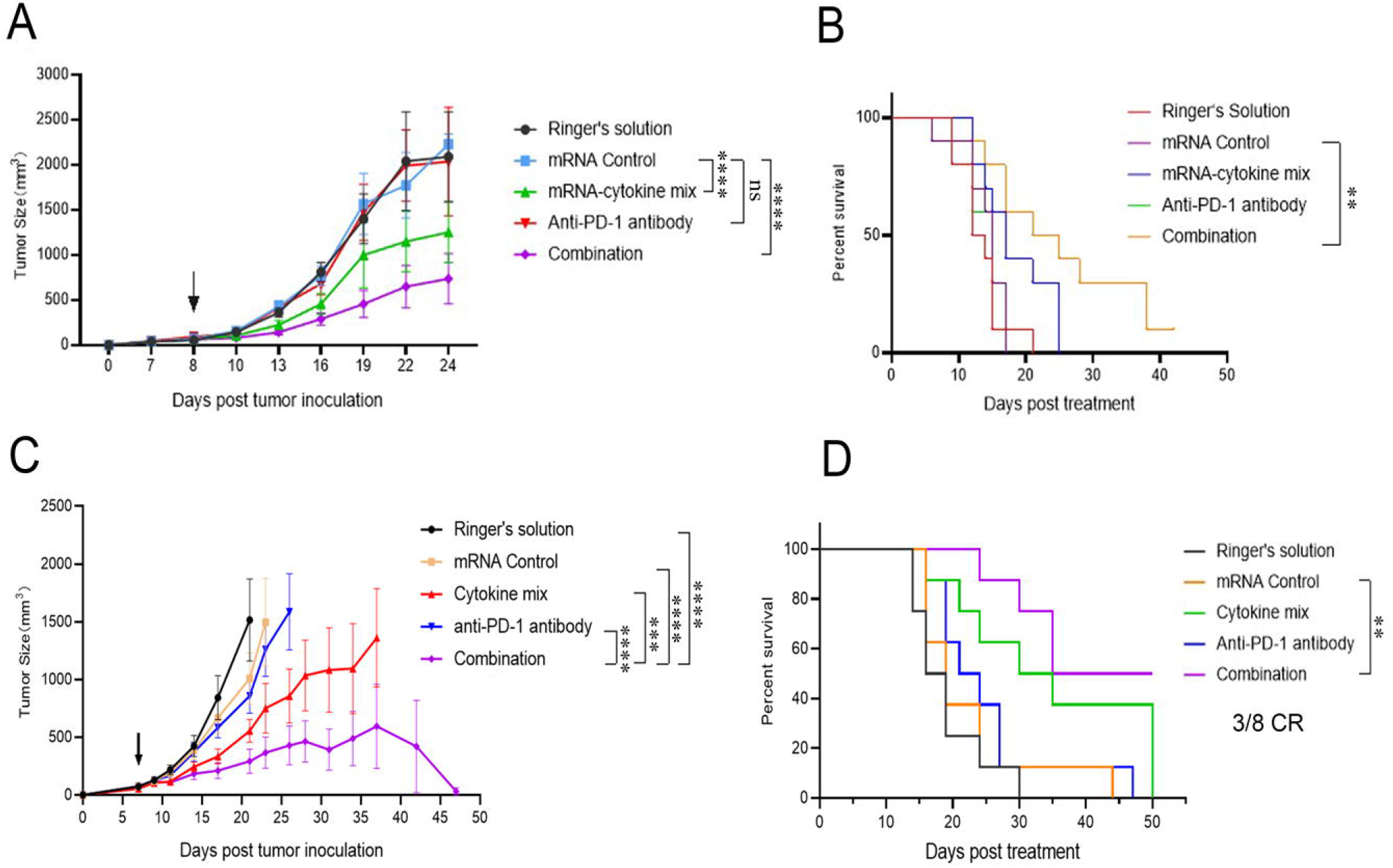
(A) C57BL/6 mice bearing B16F10 tumor were i.t. injected with indicated solution or naked C-RNA and/or i.p. injected with anti-PD-1 antibody. In vivo antitumor activity of mRNA encoding a mixture of four C-RNA cytokines with/without in combination with anti-mouse PD-1 antibody was evaluated by the tumor growth over time as well as survival curve. (B) C57BL/6 mice bearing MC38 tumor were i.t. injected with indicated solution or naked C-RNA and/or i.p. injected with anti-PD-1 antibody. In vivo antitumor activity of mRNA encoding a mixture of four C-RNA cytokines with/without in combination with anti-mouse PD-1 antibody was evaluated by the tumor growth over time as well as survival curve. Data plotted are group mean tumor volume ±SEM. For (A) (B), n=10, for (C) (D), n=10. **P<0.01, ****P<0.0001.

### Cytokines-encoding C-RNA suppressed tumor growth by promoting infiltration and activation of T cells

We proposed that cytokine C-RNA mixture modulates tumor microenvironment, especially the activation of T cells for tumor suppression. To delineate the cellular mechanism for cytokine C-RNA mixture mediated tumor growth suppression, tumor-infiltrated T cells were analyzed by flow cytometry after two doses of combination therapy (Fig.8A). Firstly, the percentage of infiltrated CD45^+^cells were analyzed, and results indicated that the treatment with C-RNA mixture elicited promotion of percentage of CD45^+^ cells; importantly, the combination treatment with C-RNA mixture and anti-PD-1 antibody led to a potent elevated percentage of CD45^+^ cells (Fig.8B), suggesting that C-RNA coordinated with anti-PD-1 antibody to facilitate the infiltration of total leukocytes into tumor. We also assessed the percentage of tumor-infiltrated T cells, as shown in Fig.8C, compared to anti-PD-1 antibody monotherapy group in which there was only 0.6% of T cells infiltrated, the combination treatment increased the percentage up to 6%. In other words, the combination treatment turned the T cell-poor tumor into T cell-inflamed tumor. The activation of recruited T cells was clarified according to the expression level of the activation marker CD69. As shown by Fig.8D., the percentage of CD69 positive T cells in total CD4 T cells was increased to 80% in C-RNA monotherapy as well as C-RNA and anti-PD-1 antibody combination therapy. However, the value was only around 60% in control or anti-PD-1 antibody monotherapy group. Likewise, it was also the case for tumor-infiltrated CD8 T cells. C-RNA monotherapy and combination treatment elevated the percentage of CD69 positive cells up to 70% and 80% respectively. Collectively, not only there were more immune cells especially T cells recruited to the tumor, but also their activation degrees were higher, thus exerting stronger anti-tumor effects.

**Fig.8.**
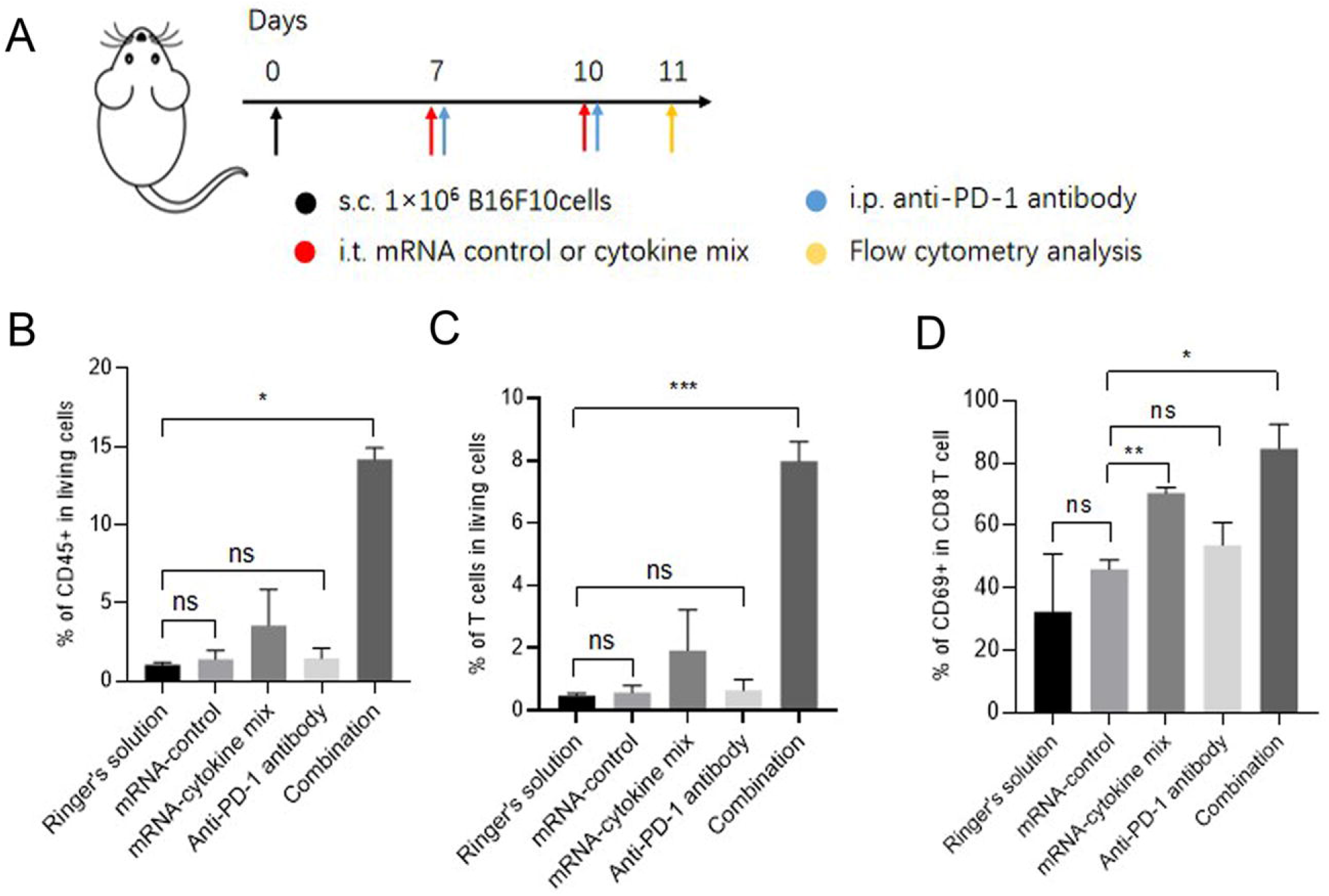
(A) Experiment timeline for treatment and analysis of B16F10 tumor-bearing mice. s.c., subcutaneous; i.t., intratumoral; i.p., intraperitoneal; (B)Percentage of CD45 positive cells in living cells of the tumor; (C)Percentage of CD69 positive cells in CD4 T cells in the tumor; (D)Percentage of CD69 positive cells in CD8 T cells in the tumor.

## Discussion

The method with PIE system of generating circular RNA in vitro has been developed as early as 1990s^17^, however, the application for protein expression by in vitro-generated circular RNA hasn’t been realized until recently. This is probably due to the fact that the notion of circular RNA mediated protein expression has just been conceived in recent years. The pioneering work from R Alexander Wesselhoeft and Daniel G Anderson *et*.*al* with a series of engineering of circular RNA paved the way for the potential applications of circular mRNA as vaccines and therapeutics^12^. However, more in vitro and in vivo evidence is required to prove this concept that circular mRNA can be applied to express varies types of proteins, and probably clinical studies are required to prove its potentiation as vaccines and therapeutics. In the current study, we developed a novel format of circular mRNA, termed C-RNA, which can be generated by PIE system and uses E29 IRES to direct protein translation. Our experimental results demonstrated that C-RNA mediated high activity of protein translation, as revealed by the high expression of EGFP. Importantly, we found that HPLC-grade RNA purifications resulted in notably higher protein expression in immunogenic sensitive A549 cell line. Due to the fact that immunogenic RNA triggers innate immunity, which may suppress protein expression and caused inflammation of human body^21^, therefore, the purifications of C-RNA by HPLC is crucial for the future development of RNA vaccines and therapeutics. Next, we evaluated whether C-RNA can direct the expression of varies types of proteins, such as secreted proteins and transmembrane proteins both in vitro and in vivo. We proved that C-RNA can direct the in vivo expression of intracellular protein luciferase in mice, via the mRNA delivery by LNP encapsulation. The LNP encapsulation of circular mRNA is similar as linear mRNA and the location of luciferase expression is mostly in the liver, this is consistent with the case of LNP mediated linear mRNA delivery^22^. Importantly, we observed that the expression of luciferase in local injection muscle lasted to 9 days, an extreme long duration for mRNA in vivo expression, demonstrating the high stability of C-RNA. Furthermore, a secreted protein, SARS-Cov-2 RBD antigen^23^, was encoded by C-RNA, and significant expression was validated in 293T cell line as well as in mice, demonstrated that C-RNA directs high expression of secreted protein both in vitro and in vivo. For the evaluation of membrane protein expression, C-RNA directed IL-15-CD8 α or IL-15-GPI membrane proteins were confirmed to be highly expressed in 293T cell line, suggesting that membrane proteins are suitable expression targets of C-RNA. Collectively, we conclude that C-RNA platform is sufficient for the expression of varies proteins, including intracellular, secreted and transmembrane proteins both in vitro and in vivo. Thereof, it is rational to prospect that C-RNA has the potential as vaccines or therapeutics by expressing varies types of proteins, such as antigens, cytokines, antibodies, transmembrane ligands, receptors, transcription factors, etc.

In 2016, a report from Karine Breckpot *et*.*al* revealed that direct intratumoral injection of naked linear mRNA mixture that encoding CD40L, constitutive active TLR4 and CD70 resulted in the activation of tumor infiltrated DCs and presented anti-tumor effects^24^. This study suggested that linear mRNA can be direct injected to tumor for protein expression. As previously described, naked DNA can be uptaken by cancer cells^19^. However, it is still unknown that if naked circular mRNA, a ring-shape molecule, can be uptaken by cancer cells and express in tumor tissue. In the current study, we provide evidence to prove that C-RNA, a new circular mRNA format, can be utilized for intratumoral injection. Firstly, our results demonstrated that naked C-RNA mediated high protein expression by direct injection to tumor. Moreover, C-RNA directed intratumoral expression was found to be more durable than the typical linear mRNA. This is of great importance for the application of mRNA as vaccines or therapeutics. The longer duration of mRNA directed protein expression may facilitate the development of prophylactic vaccines or therapeutic cancer vaccines, by increasing the in vivo production of recombinant viral antigens and eliciting higher titer neutralizing antibodies or stronger antigen-specific T cell activation; may benefit the development of mRNA monoclonal or bispecific antibodies by improving the in vivo pharmaceutical kinetics; may facilitate the development of mRNA based engineered Car-T/NK by increasing the duration of chimeric antigen receptor expression in T cells or NK cells. With Ringers’ solution as formulation, C-RNA can be delivered to tumor efficiently and conducted stronger protein expression. Collectively, for the first time, these results demonstrated that, as a new kind of mRNA, the naked C-RNA can be delivered to tumor tissue for protein expression by intratumoral administration, therefore, has the potential as an RNA platform to develop a variety of intratumoral therapeutics.

By direct injection of immunological regulatory agents to solid tumors, intratumoral immunotherapy is a promising method to modulate tumor microenvironment^25^. Several kinds of therapeutics have shown their potential for clinical applications, for example the oncolytic virus product T-VEC, which was approved by FDA in 2015 for the therapy of advanced melanoma^26^. T-VEC conducts the lysis of tumor cells and results in release of tumor-derived antigens (TDA), and simultaneously expressing and releasing GM-CSF to activate dendritic cells (DCs) and T cells to trigger immune responses to tumor antigens, finally gives rise to distant immune responses and causes a systemic anti-tumor effect. This mechanism of action is designated as *in situ* vaccination^27^, and thus results in a preferable therapeutic effect with the combination of immune checkpoint blockade agents^28^. Intratumoral injection of mRNA is an emerging research area and presents outstanding therapeutic results in recent years. The pipeline of LNP-delivered linear mRNA intratumoral injection developed by Moderna includes mRNA-2905 (IL-12)^29^ and mRNA-2752 (a mixture of IL-23, IL-36γ and OX40L)^30^. In the pre-clinical studies, these therapeutic candidates exhibited remarkable anti-tumor effects with the combination of immune checkpoint blockade agents. The basic principle of these candidates is the modulation of tumor microenvironment by boosting immune cells infiltration and activation, ultimately eliciting systemic anti-tumor responses. In the current study, the therapeutic effect of a mixture of circular mRNA that encoding cytokines including IL-15, IL-12, GM-CSF and IFN-α 2b was assessed as the proof of concept. Our results suggested that C-RNA directed high quantity expression of those cytokines *in vitro*, as detected by ELISA. Furthermore, the *in vitro* expressed cytokines were found to activate immune cells after transferring to PBMC, demonstrating that C-RNA-derived cytokines are highly active. The intratumoral administration of C-RNA mixture resulted in remarkable regression of tumor growth in the B16F10 syngeneic mouse tumor model, demonstrating that C-RNA directed local expression of cytokines in tumor exhibited significant tumor suppression effect. Moreover, mechanism studies revealed that the proportion as well as the activation degree of tumor infiltrated cytotoxic CD8 T cells were up-regulated, suggesting that the tumor microenvironment was actually remodeled by the locally expressed cytokines, and facilitated anti-PD-1 mediated immune therapy. The “cold” B16F10 tumor was turned to be “hot” after intratumoral injection of C-RNA that encoding cytokines. These results are consistent with the outcomes from previous studies of intratumoral mRNA for microenvironment modulations^30^, and suggests that C-RNA is an excellent vector to express immune-modulatory factors intratumorally for cancer therapy.

In summary, we established a novel form of circular mRNA, termed C-RNA, in which the high effective protein expression is driven by E29, an IRES element derived from echovirus 29. This kind of circular mRNA is more stable and mediates higher and more durable protein expression than linear mRNA. The HPLC-grade purification of C-RNA resulted in promotion of protein expression in immunogenic cell line, probably due to the removal of immunogenic 5’-ppp intron fragments, suggesting that C-RNA is safe for in vivo usage as potential vaccines or therapeutics. Furthermore, the successful expression of varies kinds of proteins, including intracellular proteins, secreted proteins and transmembrane proteins, we conclude that C-RNA is a universal vector for protein expression. The intratumoral expression of C-RNA is found to direct higher and more durable protein expression than typical linear mRNA. The intratumoral administration of C-RNA mixture encoding four cytokines elicited notable tumor suppression effect by boosting infiltrated T cells to facilitate immune therapy. Collectively, C-RNA is proven to be a potential platform for the development of mRNA vaccines or therapeutics, may be a preferable RNA vector due to its simplicity for manufacturing and excellent performances for protein expression. Figure Legend

